# Genomic evidence for facultative selfing in the cichlid fish *Cyphotilapia frontosa*

**DOI:** 10.64898/2026.05.13.724898

**Authors:** M. Efe Uysal, Daniela Souza-Costa, Allison Marks, Adrian Indermaur, Wolfgang Gessl, Walter Salzburger, Julia M. I. Barth

## Abstract

Organisms have evolved a remarkable diversity of reproductive strategies in response to environmental variations and selective pressures. Although most vertebrates do reproduce biparentally, rare alternative modes such as selfing (self-fertilization) and different forms of parthenogenesis exist, but remain poorly characterized. Here, we investigated an unusual reproductive event in the normally biparental cichlid fish *Cyphotilapia frontosa*, in which a female produced offspring in the absence of a male. Using whole-genome sequencing data, we analyzed whether reproduction occurred via selfing or parthenogenesis by comparing patterns of heterozygosity with those from a wild, genetically diverse *C. frontosa* family collected in Lake Tanganyika and a closely related inbred *Ctenochromis benthicola* family. The uniparental family exhibited reduced genetic diversity, elevated relatedness, and genome-wide patterns of homozygosity distinct from those expected under parthenogenesis or inbreeding, but consistent with self-fertilization. Our study provides rare genomic evidence of selfing in a vertebrate and suggests that such alternative reproductive modes may be overlooked rather than truly absent. These findings contribute to a broader understanding of how alternative reproductive strategies evolve in vertebrate lineages.

**Significance:** The overwhelming majority of vertebrates reproduce sexually, requiring a male and a female to produce genetically distinct offspring. Yet, rare alternative modes involving only a single parent such as asexual parthenogenesis (“virgin birth”) or self-fertilization challenge this paradigm. Among these, selfing is exceptionally uncommon and poorly studied in vertebrates. Here, we unveiled - based on genomic analyses - the reproductive strategy of a member of the extraordinarily diverse cichlid fish radiation in Lake Tanganyika that reproduced in captivity in the absence of a male. By comparing patterns of genome-wide heterozygosity with both wild and inbred reference families, we identified a rare case of selfing. This finding adds to the limited records of selfing in vertebrates and expands current understanding of reproductive diversity, highlighting the power of whole-genome sequencing to distinguish among alternative reproductive mechanisms.

## Introduction

Reproduction is a fundamental biological process and essential for the continuation of species through generations (Fusco & Minelli, 2023). In order to maximize their reproductive success, organisms use various reproductive strategies (Stearns, 2000; Kappeler, 2012; Shivanna & Tandon, 2014), whereby reproductive modes are typically categorized into two main groups: sexual and asexual reproduction (**Fig. 1**) (Dimijian, 2005; Fusco & Minelli, 2019). In vertebrates, the predominant mode of reproduction is sexual, or more specifically, biparental sexual (Lombardi, 1998). However, a number of cases have been reported where vertebrates reproduce uniparentally through parthenogenesis or selfing (Neaves & Baumann, 2011; Avise, 2015; Kelley et al., 2016). These relatively rare cases are noteworthy because they reveal that vertebrate reproduction is not strictly biparental, and that different reproductive strategies may have distinct consequences on genetic diversity, population growth, and evolution (Hörandl et al., 2020; Fujita et al., 2020).

**Figure 1.**
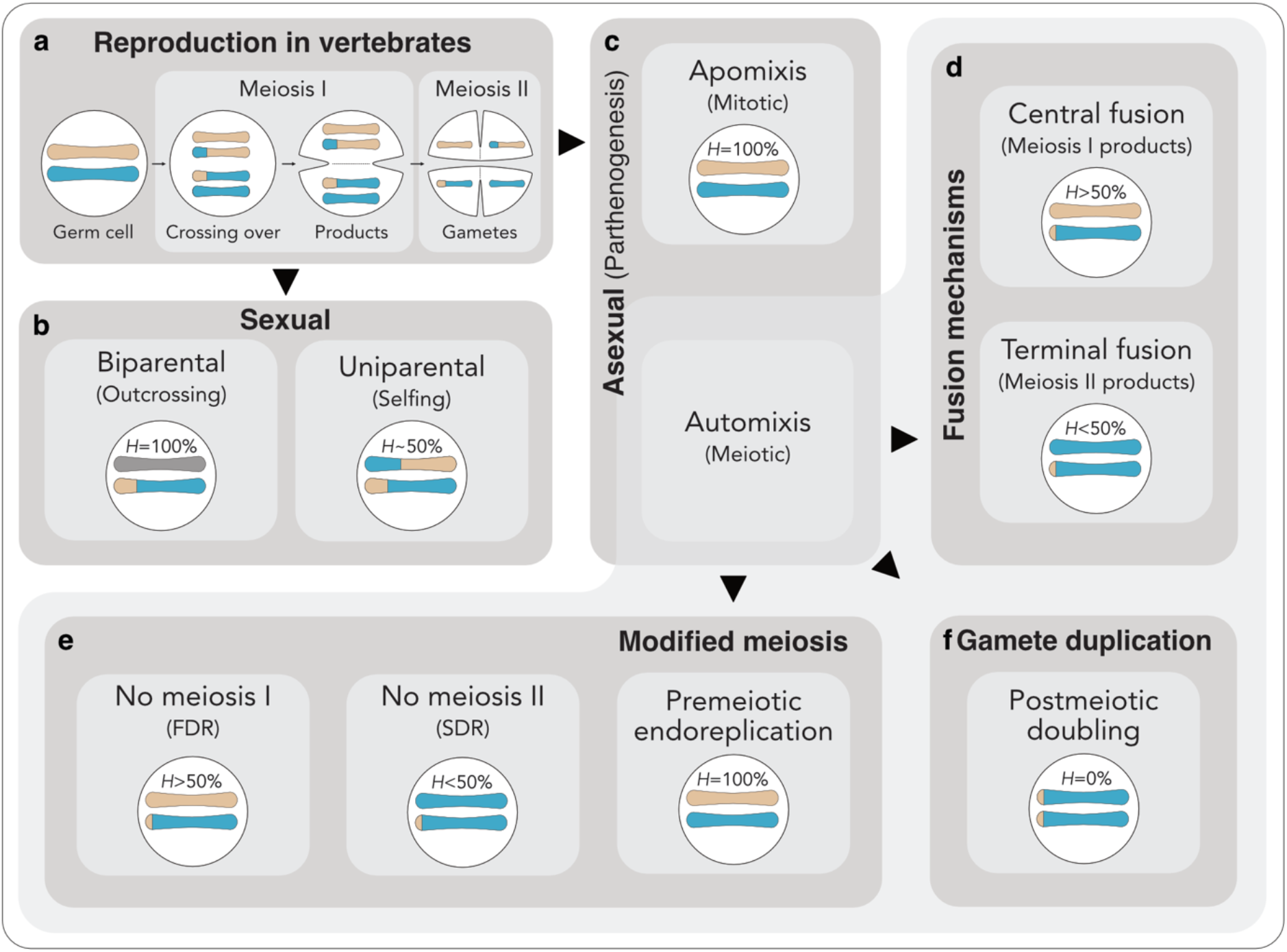
Reproductive modes and their genomic consequences in vertebrates. During canonical meiosis, DNA replication is followed by two successive divisions (meiosis I and II) to produce haploid gametes (**a**). Diploidy is restored in sexual reproduction either by fusion of gametes from two individuals (outcrossing), maintaining genome-wide heterozygosity, or by fusion of two independent gametes from the same individuum (selfing), resulting in genome-wide reduction and stochastic redistribution of heterozygosity (**b**). In asexual (parthenogenetic) reproduction, diploidy is restored without fertilization, either via mitotic divisions (apomixis) producing clonal offspring with fully retained heterozygosity, or via altered meiosis (automixis) (**c**). Automixis comprises distinct cytological mechanisms with characteristic genomic outcomes: Fusion of meiotic products (**d**) can occur after meiosis I (central fusion), reuniting homologs and retaining high heterozygosity, particularly in regions of low recombination (e.g., near centromeres), or after meiosis II (terminal fusion), where fusion of sister chromatids results in extensive homozygosity. Alternatively in automixis, diploidy can be restored through modification of meiotic division itself (**e**): Suppression of meiosis I (first division restitution, FDR) prevents homolog segregation and largely preserves heterozygosity, whereas suppression of meiosis II (second division restitution, SDR) allows homolog segregation but retains sister chromatids, resulting in partial loss of heterozygosity with recombination-dependent patterns. In premeiotic endoreplication, genome duplication prior to meiosis leads to pairing of identical copies, effectively bypassing homolog interactions and preserving heterozygosity. Finally, postmeiotic genome duplication (gamete duplication) (**f**) restores diploidy by doubling a haploid genome, resulting in complete homozygosity. Percentages display retained heterozygosity (*H*).

In biparental sexual reproduction, haploid gametes (sex cells) are produced via meiosis in each parent and fuse to restore diploidy in the zygote (**Fig. 1a**) (Mackiel & Dillon, 2021). Recombination occurs during meiosis, providing an opportunity for DNA repair and creating new combinations of alleles, thereby contributing to genetic diversity among offspring (Mirzaghaderi & Hörandl, 2016; Hegde & Crowley, 2019). Selfing is also a form of sexual reproduction, albeit with the difference that the sex cells arise from a single individuum (‘uniparental sexual reproduction’) (**Fig. 1b**) (Jarne & Auld, 2006). As an extreme form of inbreeding, selfing is expected to reduce heterozygosity (*H*) by approximately 50% per generation (Cornetti et al., 2021). Asexual parthenogenetic reproduction, on the other hand, does not involve the fusion of gametes (Gavrilov-Zimin, 2023). Instead, the embryo develops from an unfertilized egg, or in other words, without any sperm being involved, and diploidy is restored by other mechanisms (**Fig. 1c**) (Lampert, 2008). There are several types of asexual reproduction, which vary in their underlying cellular mechanisms and therefore result in different heterozygosity patterns in the offspring (Stenberg & Saura, 2009; Glémin, 2019). For example, in mitotic parthenogenesis (apomixis), meiosis is absent, and the offspring are genetically identical (clonal) to the parent, thereby fully retaining maternal heterozygosity patterns (Mirzaghaderi & Hörandl, 2016). Yet, not all forms of clonal reproduction are completely male-independent, with some – such as gynogenesis – requiring a sperm cell to activate the process albeit without genetically contributing to the offspring (Avise, 2015). In automixis with central or terminal fusion, both meiotic divisions are completed and diploidy is restored by the fusion of meiotic products derived from meiosis I or meiosis II, which, in oogenesis, correspond to fusion with the first or second polar body, respectively (**Fig. 1d**) (Lampert, 2008). As a consequence, heterozygosity is either lost or retained, depending on the meiotic products that are fused (Card et al., 2021). Other parthenogenetic strategies restore diploidy by modifying meiosis (**Fig. 1e**). This can involve genome duplication prior to meiosis (premeiotic endoreplication), allowing pairing to occur between identical sister chromosomes rather than homologous chromosomes and ensuring the full retention of maternal heterozygosity in the offspring. Alternatively, diploidy can be restored through suppression of meiotic divisions: suppression of meiosis I prevents homolog segregation and preserves heterozygosity, whereas suppression of meiosis II allows homolog segregation but retains sister chromatids, resulting in partial, recombination-dependent loss of heterozygosity with spatial structuring along chromosomes (Lutes et al., 2010; Smith et al., 2019). Finally, post-meiotic doubling (gamete duplication) closely resembles terminal fusion automixis with a substantial loss of maternal heterozygosity (Stenberg & Saura, 2009). Similar to gynogenesis, some meiotic modifications require male contribution. In hybridogenesis, maternal meiosis produces haploid gametes that are fertilized by sperm, but the paternal chromosomes are discarded during oogenesis (Avise, 2015).

Despite asexual parthenogenetic reproduction being present in many invertebrates (Normark 2003; Jaron et al., 2021), in vertebrates obligate parthenogenesis exclusively occurs in squamate lizards and snakes (Fujita et al., 2020). Examples of facultative parthenogenetic reproduction in vertebrates include the king cobra (*Ophiophagus hannah*) and the common smooth-hound shark (*Mustelus mustelus*), both inferred to reproduce via automixis with terminal fusion (Card et al., 2021; Esposito et al., 2024), and several geckos, which can reproduce via premeiotic endoreplication (Dedukh et al., 2022). In contrast, selfing seems to be exceedingly rare in vertebrates and is well-established only in the mangrove killifish *Kryptolebias marmoratus* (Kelley et al., 2016). In cichlid fishes, biparental sexual reproduction has been described as the standard mode (Sefc 2011), with selfing occurring under certain conditions as shown in *Benitochromis nigrodorsalis* (Böhne et al., 2023), and a cichlid hybrid (Svensson et al., 2016), but its prevalence and stability remain unclear.

The analysis of whole-genome sequencing (WGS) data is a powerful approach for distinguishing between different reproductive modes, on the ground that each reproductive strategy is expected to leave characteristic patterns of retained heterozygosity (Fig. 1). Moreover, in contrast to analyses based on a limited number of genetic markers, WGS data in combination with a chromosome-level reference assembly enable the identification of recombination-associated patterns along chromosomes, providing a diagnostic tool to differentiate reproductive strategies. In particular, centromeric regions are informative, as recombination rates are typically reduced near centromeres and elevated toward telomeres, breaking up linkage between alleles (Haenel et al., 2018; Sardell et al., 2018). For example, in automixis, centromere-proximal regions are expected to exhibit a consistent state within an offspring across chromosomes (heterozygosity retention under central fusion and loss under terminal fusion), whereas in selfing, retention patterns are mainly random because offspring inherit the gametes produced in independent meiotic events (Engelstädter 2017; Böhne et al., 2023). As similar homozygosity patterns may also arise from extended inbreeding, comparing genomic signatures across reproductive strategies in families with varying outcrossing and inbreeding levels is essential to disentangle their underlying mechanisms (Zeitler & Gilbert 2024).

In this study, we investigated an alternative reproductive strategy in the maternal mouthbrooding cichlid fish *Cyphotilapia frontosa* (*Cy. frontosa*), a member of the extraordinarily diverse Lake Tanganyika cichlid radiation exhibiting a remarkable range of life-history traits (Kuwamura 1986; Sefc 2011). *Cy. frontosa* reproduces under natural conditions through biparental sexual reproduction (Kuwamura 1997), yet in our case, a captive female reproduced without genetic contribution from a male. Using WGS data from three families – two biparental sexually reproducing families (one genetically diverse *Cy. frontosa* family from the wild and one inbred aquarium family of the closely related species *Ctenochromis benthicola*) and the putatively uniparental *Cy. frontosa* family – we compared relatedness among family members, tested for paternal genetic contribution, and quantified genome-wide patterns of heterozygosity retention. By mapping genomic data to a chromosome-level reference assembly we further examined heterozygosity states within centromeric regions to uniquely determine the specific reproductive strategy of *Cy. frontosa*.

## Results

### Genome-wide variation and relatedness patterns distinguish outbred, inbred, and uniparental reproductive lineages

To assess WGS data quality and patterns of inter- and intra species divergence and relatedness among families, we analyzed per-sample variant metrics, genome-wide variation, and identity-by-descent (IBD) across the two sexually reproducing families and the focal uniparental family. Variant metrics indicated high sequencing quality (transition/transversion ratio (Ts/Tv) = 2.28 ± 0.02; Table S1). The heterozygous-to-homozygous alternate ratio (het/hom alt) mirrored differences in genetic diversity, relatedness, and population origin across the three families. In the wild *Cy. frontosa* family, the putatively unrelated male exhibited the highest ratio (0.86), consistent with increased heterozygosity. The female, likely originating from a local breeding cohort, showed a lower ratio (0.65), while the five offspring had a mean ratio of 0.51± 0.07. In contrast, the aquarium-bred *Ct. benthicola* family showed uniformly low values (0.16 ± 0.004), a clear inbreeding signature. The uniparental *Cy. frontosa* family exhibited intermediate values (0.53), with reduced values in the offspring (0.24 ± 0.04), consistent with reduced heterozygosity, a possible signature of parthenogenesis or selfing.

A principal component analysis (PCA) of genome-wide variation showed that the first principal component axis (PC1; 68.24%) separated *Ct. benthicola* from both *Cy. frontosa* families (Fig. S1), while PC2 (18.66%) distinguished the wild and uniparental *Cy. frontosa* families. Within the wild family, the male (specimen ID: 04G6) clustered separately from the female and offspring, while the latter formed two partially overlapping subgroups: one containing the female and two offspring (04H1 and 04H4), and the other including the remaining three offspring (04G8, 04H2, and 04H5). In the uniparental family, one offspring (AM03) appeared genetically distinct; however, its high rate of missing data and elevated singleton count (Table S1) suggests this pattern likely reflects lower sequencing quality and genotyping errors rather than true biological divergence.

Identity-by-descent (IBD) analysis quantified the proportion of shared genome (PI_HAT) as 0.5 × Z1 + Z2, where Z1 and Z2 represent the probabilities of sharing one or two alleles by descent, respectively. In the wild *Cy. frontosa* family, siblings (PI_HAT 0.65 ± 0.04) and parent–offspring pairs (0.73 ± 0.05) showed moderately elevated allele sharing relative to outbred expectations (0.5), consistent with relatedness within a local breeding cohort (Table S2). The parental pair also exhibited elevated allele sharing (0.55), suggesting either relatedness of male and female or that linkage-filtering thresholds were too lenient. The aquarium-bred *Ct. benthicola* family showed very high genome-wide allele sharing (full siblings: 0.93 ± 0.01; parent-offspring pairs 0.90 ± 0.002; parental pair 0.74), indicating high relatedness. The uniparental *Cy. frontosa* family displayed a distinct pattern with high sibling PI_HAT (0.77 ± 0.06) driven by elevated Z2 values (0.76 ± 0.06) and low Z1 values (0.03 ± 0.07, (Fig. S2) indicating offspring share both alleles at the majority of loci. Mother-offspring comparisons were also unusually high (0.89 ± 0.01). Together these patterns are consistent with uniparental reproduction in the absence of male contribution, producing near-clonal offspring.

### High genotype sharing and absence of paternal genetic contribution support selfing or uniparental parthenogenetic reproduction

We tested for evidence of clonality and paternal contribution by calculating the proportion of shared genotypes (*M*_xy_) and the proportion of non-Mendelian genotypes within families, respectively. The wild *Cy. frontosa* family showed the lowest within-family similarity with *M*_xy_ 0.76 ± 0.02 between siblings, *M*_xy_ 0.78 ± 0.04 for parent-offspring comparisons, and *M*_xy_ 0.65 between parents (**Fig. 2a**; Table S3), matching expectations for an outbred, naturally mating population. In contrast, the *Ct. benthicola* family had the highest genotype similarity with *M*_xy_ 0.95 ± 0.005 across siblings, *M*_xy_ 0.92 ± 0.002 between parents and offspring, and high similarity (*M*_xy_ 0.87) also between parents (**Fig. 2a**; Table S3), consistent with an inbred lineage with very low genetic diversity. The uniparental *Cy. frontosa* family, suspected to have reproduced without paternal contribution, showed high *M*_xy_ values (0.9 ± 0.015 for siblings; 0.91 ± 0.012 for parent–offspring; **Fig. 2a**; Table S3). These values are lower than expected for clonal reproduction (*M*_xy_ ∼1.0) and also below the proportion of shared genotypes observed in the inbred *Ct. benthicola* family, indicating that technical issues, such as genotyping errors, are unlikely to explain downwardly biased estimates. Instead, their *M*_xy_ is consistent with alternative reproductive strategies like selfing or uniparental meiotic parthenogenesis, which can produce offspring ranging from partially to fully identical to the mother, depending on the underlying cytological mechanism (**Fig. 1 d-f**).

**Figure 2.**
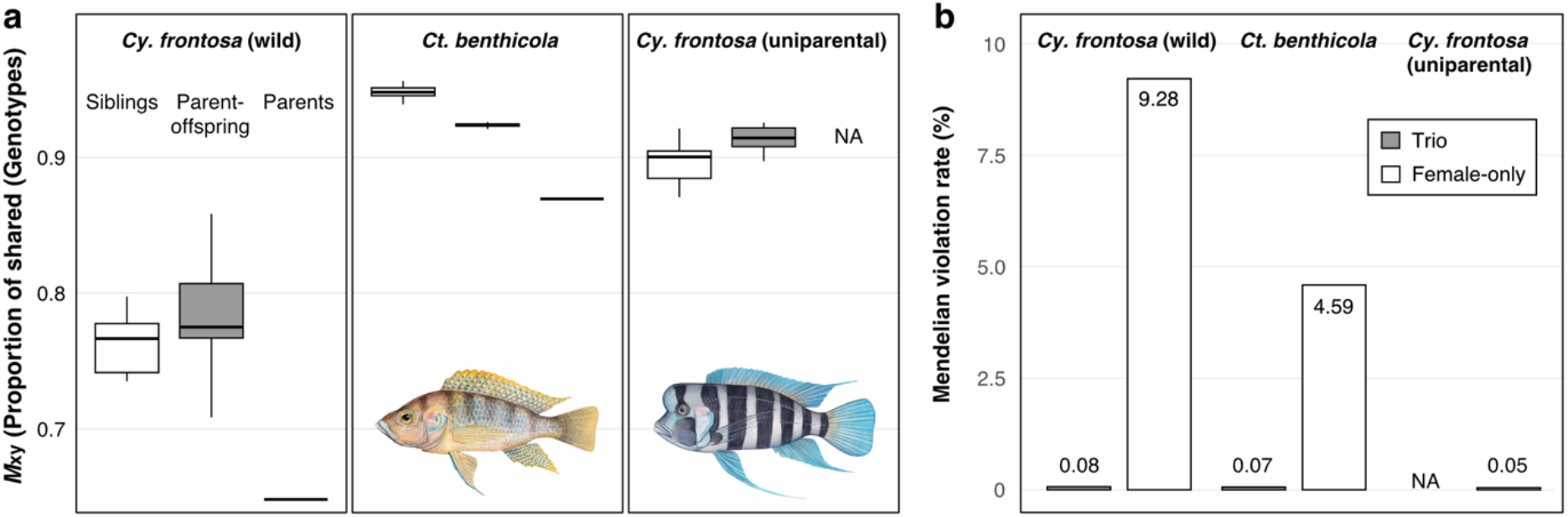
Proportion of shared genotypes (*M*_xy_) and non-Mendelian genotypes. **(a)** Proportion of shared genotypes (*M*_xy_) among siblings and parent-offspring pairs for all families, and between parents of wild *Cy. frontosa* and *Ct. benthicola*. Boxplots show the interquartile range (25th–75th percentiles), the thick line indicates the median, and whiskers extend to 1.5× the interquartile range. Images depict the two species. **(b)** Mendelian violation rate in trio (male, female, offspring) and female-only comparisons. NA indicates unavailable male estimates for the uniparental family.

Furthermore, Mendelian inconsistency rates were low across all three families (wild *Cy. frontosa* 0.08%; *Ct. benthicola* 0.07%; uniparental *Cy. frontosa* 0.05%; **Fig. 2b**; Table S4), consistent with residual genotyping errors or *de novo* mutations. In contrast, if a male had contributed genetically to the uniparental *Cy. frontosa* family, we would have expected substantially higher apparent Mendelian violation rates, similar to those observed when paternal genotypes are ignored in the sexual families (wild *Cy. frontosa* 9.28%; *Ct. benthicola* 4.59%; **Fig. 2b**; Table S4). This ∼200-fold difference strongly supports the absence of paternal genetic contribution in the uniparental *Cy. frontosa* offspring. For both tests, analyses with the non-LD pruned dataset (2,690,649 SNPs) yielded similar results (Table S4).

### Retained heterozygosity distributions identify selfing as the alternative reproductive mode in *Cy. frontosa*

To further assess the underlying reproductive mode of the uniparental family, we inferred the number and percentage of retained heterozygous sites in the offspring as well as retention pattern along and across chromosomes, and within centromeric regions. Accordingly, we restricted the analyses to sites where the mother is heterozygous in the uniparental family, or where at least one parent is heterozygous in the biparental families and the offspring had no missing data (number of informative sites). This number differed greatly among the three families, reflecting differences in genetic diversity within each family, but was relatively consistent across offspring within each family (Table S5). The genome-wide retained heterozygosity was around 50% for all three families (49.42 ± 4.38%). The wild biparental *Cy. frontosa* family exhibited large variation among offspring (48.98 ± 5.12%), whereas the biparental inbred *Ct. benthicola* family had a narrow range (50.05 ± 1.47%), indicating lower variability among the offspring. Although outcrossing is expected to maintain heterozygosity genome-wide (**Fig.1b**), the ∼50% heterozygosity observed at informative sites in biparental families arises from Mendelian segregation at loci where at least one parent is heterozygous. In the uniparental *Cy. frontosa* family, heterozygosity retention ranged from 49.26 ± 6.83%), with one offspring (AM08) showing notably lower retention compared to the remaining offspring, which may reflect differences in recombination patterns (**Fig. 3c**).

**Fig. 3.**
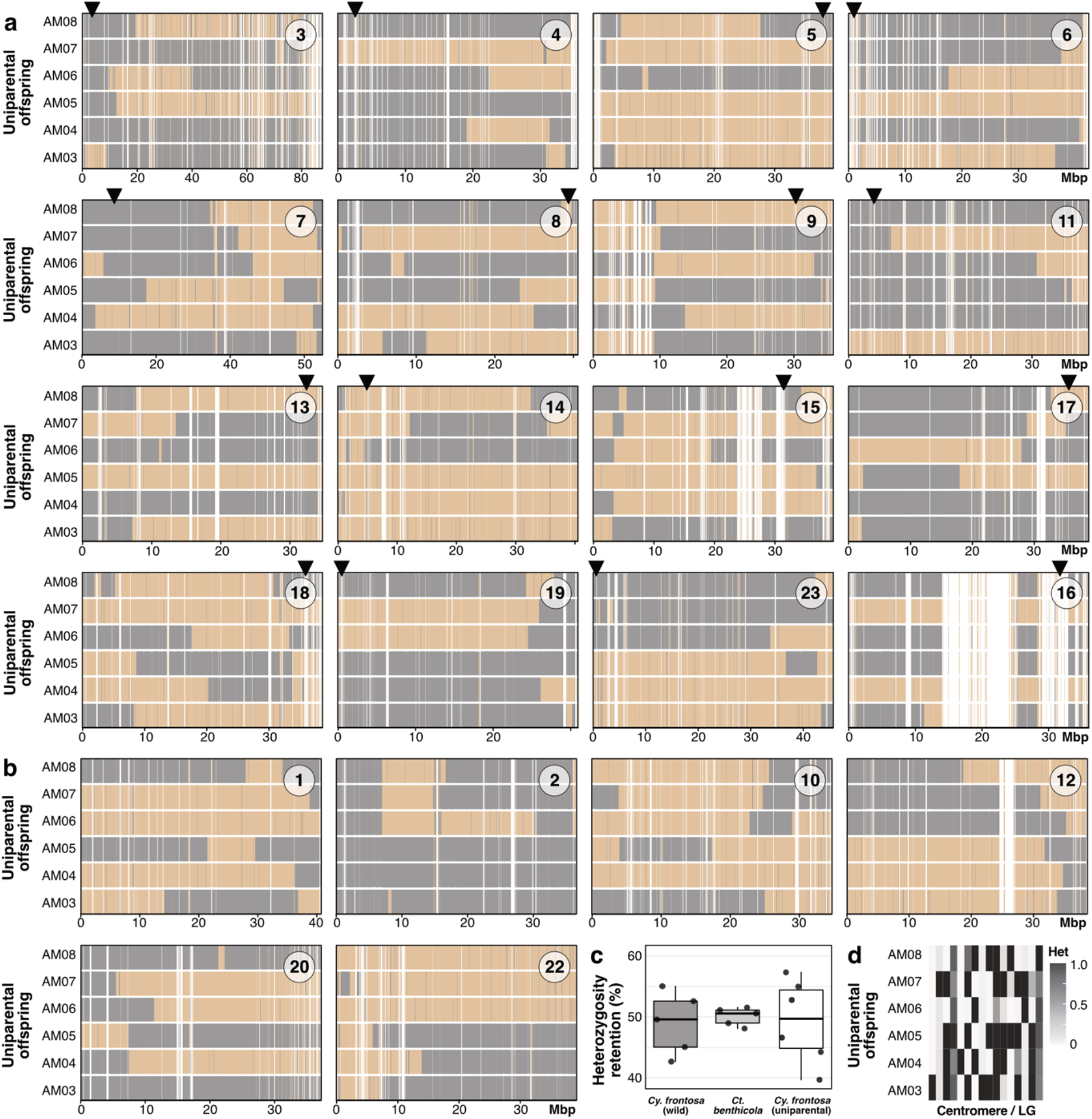
Genome-wide heterozygosity retention. **(a)** Retention profiles across 16 chromosomes with known centromeres. Vertical lines represent variant sites at their genomic position for all offspring of the uniparental *Cy. frontosa* family (AM03-08). Homozygous sites are indicated in gray, heterozygous sites in gold, white is either homozygous in the parent or missing data. Numbers display linkage groups (LGs) based on the *Oreochromis niloticus* reference (NCBI GCF_001858045.2). Centromere positions are indicated with black triangles. **(b)** Retention profiles of chromosomes without known centromere position. **(c)** Boxplot of genome-wide heterozygosity retention for all three families. Points represent individual offspring, boxes show the interquartile range (25th–75th percentiles), the thick line indicates the median, and whiskers extend to the most extreme values within 1.5× the interquartile range. **(d)** Heatmap of heterozygosity retention rates within centromeric regions of max. 2 Mb across the offspring of the uniparental *Cy. frontosa* family.

At sites with loss of heterozygosity, genotypes were either homozygous for the reference or the alternative allele. In both biparental families, homozygous reference calls were more frequent (mean 28.14 ± 1.82) than alternative calls (22.35 ± 1.82), reflecting reference bias. In the uniparental *Cy. frontosa* family, losses were evenly split (25.28 ± 3.38 vs. 25.46 ± 3.46; Table S6) as restricting analysis to maternally heterozygous sites eliminated this bias. Genome-wide heterozygosity retention in the uniparental *Cy. frontosa* family fell within a similar range to that observed the biparental families, suggesting that reproductive modes such as mitotic parthenogenesis or premeiotic endoreplication (100% retained heterozygosity), as well as gamete duplication (substantial loss of heterozygosity), are unlikely. Instead, the observed heterozygosity retention rates are more consistent with selfing or suppression of meiosis II as the underlying mechanism of uniparental reproduction.

To distinguish between these scenarios, heterozygosity retention/loss profiles were generated for each chromosome for the uniparental *Cy. frontosa* family and plotted together with the available centromere positions to examine chromosomal patterns of retention (**Fig 3a, b**; Table S6). The profiles reveal substantial variation in heterozygosity among offspring and across chromosomes. While in some chromosomes for most offspring close-to-complete heterozygosity retention was observed (e.g., LG5, 14; **Fig. 3a**), others exhibit close-to-complete loss of heterozygosity (e.g., LG2, 4; **Fig. 3a, b**). This indicates that heterozygous sites are not uniformly distributed along and among chromosomes, supporting a recombination-dependent reproductive mechanism.

Moreover, runs of homozygosity (ROH) distributions showed a clear correspondence between ROH structure and heterozygosity retention patterns in the uniparental *Cy. frontosa* family (Figure S3). To place these patterns in a broader context, we also analysed ROH distributions in the biparental families. Compared to the uniparental family, the wild *Cy. frontosa* family exhibited mostly shorter and sparse ROHs, reflecting regular outcrossing with sex-specific differences likely driven by variation in recombination or demographic history. The inbred *Ct. benthicola* family showed extensive ROH tracts across individuals, consistent with strong genome-wide homozygosity (Fig. S3).

Rates of heterozygosity retention and loss within centromeric regions differed strongly across chromosomes within each offspring. Among chromosomes with available centromere estimates (16 out of 22), centromeric heterozygosity retention ranged from 0.2% to 100%, indicating that some centromeres exhibited close-to-complete retention, whereas others exhibited close-to-complete loss (Fig. 3d; Table S7). The offspring differed in their overall centromeric retention distributions. For example, AM05 showed high retention (≥90%) in 9 out of 16 centromeric regions, whereas AM06 showed low centromeric retention (≤10%) in 10 out of 16 regions. The remaining offspring (AM03, AM04, AM07, AM08) displayed a mixed pattern, with both high and low centromeric retention rates across chromosomes. The mean heterozygosity retention rate (and mean number of SNPs) across centromeric regions for each offspring were 60.2% (307) in AM03, 49.7% (346) in AM04, 62.4% (344) in AM05, 27.9% (348) in AM06, 55.0% (346) in AM07, and 42.9% (345) in AM08 (Table S7). Analyses using 0.5 and 1.5 Mb half-widths produced the same overall pattern as the 1 Mb analysis (Fig. S4).

Centromeric regions did not show consistent heterozygosity retention or loss across chromosomes in any of the offspring. Instead, a rather random distribution pattern was observed. This contrasts with the expectation of close-to-complete retention or loss of heterozygosity across chromosomes under central- or terminal-fusion automixis, respectively, and under suppression of meiosis II. Together with the genome-wide heterozygosity retention rate in the uniparental *Cy. frontosa* family (39.68-57.31%), our results support selfing as the most likely mode of reproduction in the uniparental *Cy. frontosa* family.

## Discussion

Self-fertilization (selfing) in vertebrates is exceptionally rare compared to plants and some invertebrates. In this study, we identified the uncommon case of uniparental reproduction via selfing in the cichlid fish *Cy. frontosa* using whole-genome sequencing data by (i) assessing relatedness and differences in genetic diversity between the uniparental family and two biparental wild and inbred families, (ii) excluding possible paternal contribution, and (iii) pinpointing the underlying reproductive mechanism from genomic patterns of retained heterozygosity in the uniparental offspring.

The uniparental mother in the *Cy. frontosa* family was kept in long-term isolation as a single individual of this species in an aquarium and reproduced without a male. Consistent with this, high genotype sharing among offspring and low Mendelian violation rates indicate the absence of paternal genetic contribution. This interpretation is further supported by comparison with the biparental reference families, as the uniparental family showed distinct genomic signatures in both genetic diversity and genotype sharing for parents and parent-offspring comparisons. Likewise, identity-by-descent proportions were distinct with unusually high sibling sharing driven by elevated Z2 values compared to wild and inbred families. These observations eliminate biparental reproduction, including the high genome-wide similarity expected under biparental inbreeding, as well as male-dependent forms of parthenogenesis, such as gynogenesis and hybridogenesis.

Genome-wide heterozygosity retention in the uniparental offspring was approximately 50%, with chromosome-scale retention profiles characterized by alternating regions of retained and lost heterozygosity, also reflected in alternating runs of homozygosity (ROH), which were only partially shared among offspring. These patterns were in stark contrast to the sparse ROHs observed in the wild biparental family and the extensive genome-wide homozygosity of the inbred family. The observed patterns and the level of heterozygosity retention excludes mitotic (apomictic) parthenogenesis and premeiotic endoreplication, both of which would produce offspring genetically identical to the mother with near-complete heterozygosity retention aside from genotyping errors and *de novo* mutations. It also excludes postmeiotic genome doubling (gamete duplication), which would result in complete homozygosity and thus no retained heterozygosity (Böhne et al., 2023). Instead, the mosaic heterozygosity retention patterns observed in the uniparental *Cy. frontosa* offspring are more consistent with alternative processes involving meiosis and segregation. Notably, offspring AM08 shows the greatest deviation from the median heterozygosity retention observed in offspring, which may be explained by variation in realized recombination and segregation.

Among the alternative meiotic (automictic) modes, distinct cytological mechanisms predict retained heterozygosity, especially around centromeres. Central fusion is expected to retain high heterozygosity near centromeres, with homozygosity increasing with distance from the centromere, whereas terminal fusion leads to extensive homozygosity at centromeres, with heterozygosity increasing with distance away from it (Svendsen et al., 2015). In the uniparental *Cy. frontosa* offspring, heterozygosity retention in centromere-proximal regions is inconsistent and highly variable across chromosomes and offspring, with no uniform state across centromeres and instead a mixture of near-complete retention and near-complete loss. This variability is difficult to align with either central or terminal fusion acting uniformly genome-wide.

Alternative automictic mechanisms involving modified meiotic divisions (first or second division restitution) also do not match our observations. First division restitution (no meiosis I) largely preserves heterozygosity since homologs do not separate. In contrast, in second division restitution, meiosis II fails and sister chromatids do not separate after recombination in meiosis I, resulting in loss of heterozygosity where no recombination occurred, which is often in proximity to centromeres (Cuenca et al., 2015). Therefore, the combination of genome-wide heterozygosity retention of ∼50% and the stochastic distribution of heterozygous and homozygous regions across centromeric regions within each offspring are best explained by selfing, involving independent segregation events.

We would like to note, however, that methodical choices may influence local estimates of heterozygosity retention. In particular, sequences were mapped to the chromosome-scale assembly of *Oreochromis niloticus* (Conte et al., 2019), because of its accessibility of centromeric regions. The karyotype of *O. niloticus* comprises 22 chromosome pairs (2n = 44). As chromosome numbers in African cichlids are relatively stable, with most species possessing 22 chromosome pairs (Salzburger 2018), this provides a reasonable reference choice. However, it can not be excluded that the species investigated here differ from this number, potentially leading to differences in chromosome assignment of mapped reads, an increased proportion of unmapped reads, and reference bias. This may affect heterozygosity estimates in either direction: underestimated heterozygosity may result from unmapped reads, whereas inflated estimates can occur if reads from duplicated regions, paralogs, or structurally rearranged regions map to the same position in the reference assembly. Centromeric regions are especially prone to such bias due to their repetitive composition. Yet, we mitigated this potential impact by applying conservative quality filtering thresholds and excluding repetitive and low-mappability regions entirely. Nevertheless, the number of retained SNPs suitable for heterozygosity inference remained sufficient. Moreover, inferred centromere positions were not available for every chromosome, and the predominantly acrocentric karyotype of cichlids (Conte et al., 2019) may further contribute to low variant density in centromeric regions. These factors might affect absolute heterozygosity retention values; however, our inference of selfing as the reproductive mode relies primarily on chromosome-scale patterns and their inconsistency across 16 centromeric regions in six offspring, together with the genome-wide heterozygosity retention rate, and is therefore considered well supported.

The regular mode of sexual reproduction is biparental in vertebrates and is also predominant for cichlids (Sefc 2011) as well as the here investigated species *Cy. frontosa* (Kuwamura 1997). The percentage of heterozygous sites (Table S1) was higher in the wild *Cy. frontosa* parental individuals (20.10 ± 2.85%) and the uniparental female (15.42%) than in the inbred *Ct. benthicola* parents (5.71 ± 0.08%), supporting that biparental sexual reproduction is the predominant reproductive mode in *Cy. frontosa*, with selfing representing a facultative mode under certain environmental conditions or reproductive constraints (Lamy et al., 2011). There are a few reported cases of selfing in fishes, specifically, in the mangrove killifish *Kryptolebias marmoratus* and *K. hermaphroditus*, and in the cichlid fish *Benitochromis nigrodorsalis* (Harrington, 1961; Kanamori et al., 2016; Böhne et al., 2023). In natural populations of *K. marmoratus*, males are rare and many individuals are simultaneous hermaphrodites that reproduce via selfing (De Mitcheson & Liu, 2008; Tatarenkov et al., 2012). In cichlids, sex determination is controlled by highly dynamic sex chromosomes (El Taher et al., 2021), and sex change can occur in some species (Gammerdinger & Kocher, 2018; Oldfield, 2005). However, whether *Cy. frontosa* is capable of sex change during its lifetime is so far unknown. Moreover, Svensson et al., 2016 reported a functional hermaphrodite first-generation hybrid capable of selfing, obtained by crossing two sexually reproducing *Pundamilia* species. Therefore, selfing may be possible in cichlids under certain conditions. Notably, natural hybridization has been observed along a narrow hybrid zone between the northerly distributed *Cy. frontosa* and its southern sister species *Cy. gibberosa* (Jackson et al. 2024). Yet, our uniparental *Cy. frontosa* female shows heterozygosity estimates comparable to those of the wild parents, originating from outside the hybrid zone, making a hybrid origin unlikely as elevated heterozygosity would be expected under recent hybridization. By using selfing as an adaptive strategy, organisms can assure reproduction when males are unavailable. In contrast to asexual reproduction, selfing may also be more efficient to purge deleterious recessive mutations (Arunkumar et al., 2015; Klemp et al., 2026). Facultative selfing is therefore expected to be favoured in systems where mating opportunities are unpredictable or spatially constrained. In cichlids, such conditions may be influenced by life-history traits and temporal or spatial environmental heterogeneity shaping reproductive opportunities. *Cy. frontosa* is one of the larger Lake Tanganyika cichlids, reaching a total length of 35 cm, and inhabits deeper rocky habitats where it typically occurs in social groups consisting of a dominant male and multiple females (Konings 2019). Such a social structure, combined with spatial segregation in patchily distributed habitats in the northern part of the lake, may lead to periods of limited mate access for some individuals. In addition, *Cy. frontosa* is a maternal mouthbrooder (Kuwamura 1986; 1997) with a long brooding period of approximately 40 days and low fecundity (Konings 2019; Riehl 2001), increasing costs of reproductive failure when mating opportunities are rare or unpredictable. Under these conditions, facultative selfing could represent an alternative reproductive strategy that ensures reproductive output when males are unavailable, particularly in isolated or low-density habitats. Beyond these short-term fitness benefits, reproductive flexibility may also have longer-term evolutionary consequences. Given that variation in life-history traits is discussed to contribute to diversification in adaptive radiations (De-Kayne et al., 2025), our results suggest that such reproductive flexibility may constitute an additional mechanism facilitating rapid speciation. Future studies should therefore address two complementary questions: under which ecological and social conditions – such as low population density, reduced habitat connectivity, or skewed sex ratios – alternative reproductive modes are expressed, and whether such flexibility ultimately promotes or constrains diversification in adaptive radiations.

## Materials and Methods

### Sample collection

The *Cy. frontosa* female was kept by AM as the sole individual of this species in an aquarium for nine years before producing six offspring in 2022. The tank also contained five clown loaches (*Chromobotia macracanthus*) and one pleco (*Ancistrus sp*.), all belonging to phylogenetically distant taxa and orders distinct from Cichliformes. The wild *Cyphotilapia frontosa* family was collected in May 2021 at Lake Tanganyika, Kigoma, Tanzania. The *Ctenochromis benthicola* family was bred in the aquarium facility of the University of Graz, Austria and sampled in June 2020 (note that this species is also known as *Trematochromis benthicola*, and that we follow the nomenclature of Ronco et al., 2020 here). For all specimens, tissue samples were taken by fin-clipping and immediately stored in 96% ethanol. Sampling was done under the research permits issued by the relevant authority in the United Republic of Tanzania, and the Swiss cantonal veterinary office (permit no. 2317).

### Whole-genome sequencing

Genomic DNA was extracted from fin clips using the E.Z.N.A. Tissue Kit (Omega Bio-Tek, United States). Library preparation and sequencing was performed at the Genomics Facility Basel (Switzerland) using mechanical shearing (Covaris, 350-600 bp insert size) and the KAPAHyperPrep PCR-free Kit (Roche, Switzerland). Paired-end sequence data of 150 bp was generated on an Illumina NovaSeq 6000.

### Quality trimming and filtering

Quality of demultiplexed sequencing reads was assessed using FastQC v0.11.8 (Andrews, 120 2010). All reads were trimmed of adapter sequences, poly-G bases >10, unknown bases (N) >10, or reads with length <15 using fastp v0.21.0 and v0.23.2 (Chen et al., 2018). Lower quality reads were further trimmed to remove reads with N >4, length <40 or 60, low quality bases <20 or 26 in windows of three bases at the beginning and end of reads, and reads with an average quality <28.

### Read mapping

Reads were mapped to the NCBI RefSeq assembly of *Oreochromis niloticus* (Nile tilapia, NCBI accession GCF_001858045.2) (Conte et al., 2019) using BWA MEM2 v.2.2.1 (Vasimuddin et al., 2019), resulting files were sorted and indexed using SAMtools v.1.12 or v1.20 (Li et al., 2009), duplicates were marked, and mapping files from multiple sequencing lanes were merged with Picard-tools v2.25.5 or v3.0.0 (https://broadinstitute.github.io/picard/). Mapping statistics were assessed with SAMtools, Picard-tools’ CollectInsertSizeMetrics, and Mosdepth v.0.3.1 or v.0.3.11 (Pedersen and Quinlan, 2018).

### Masks for unreliable regions

Masks were produced to exclude duplicated and low-complexity regions of the assembly where read mapping might be ambiguous. We used the SNPable pipeline (https://lh3lh3.users.sourceforge.net/snpable.shtml), splitting the genome into overlapping 150 bp regions and applying a threshold of a minimum of 95 out of 100 reads mapping correctly. Additionally, we identified repetitive regions with RED v0.0.2 (Girgis, 2015). Masked regions were recorded in BED format and combined into a single mask file using BEDTools v.2.27.1 (Quinlan and Hall, 2010).

### Variant calling and quality filtering

Variants from reads mapped to the Nile tilapia assembly were called with GATK v.4.2.0.0 or 4.5.0.0 (McKenna et al., 2010) focusing on those reads mapped to the 22 autosomal and the mitochondrial scaffold. GATK’s HaplotypeCaller tool was used in GVCF mode excluding the pcr-indel model to write a GVCF file per sample, followed by per-chromosome joint calling with the GenomicsDBImport and GenotypeGVCFs tools under the new allele frequency model, excluding the annotations for excess heterozygosity and inbreeding coefficient, and including non-variant-sites. The resulting VCF files were hard quality-filtered using GATK’s VariantFiltration tool and BCFtools v.1.19 (Li, 2011), including sites that matched all of the following criteria: The coordinates of the site were not included in the mask file, the Phred-scaled *p*-value of Fisher’s exact test for strand bias was not greater than 40, the symmetric odds ratio test statistic was not greater than 3, the root mean square mapping quality was at least 40, the Mann-Whitney-Wilcoxon rank sum test statistic for mapping bias between reference and alternative alleles was at least -12.5 and not greater than 5.0, the quality score normalized by allele depth was at least 2, the Mann-Whitney-Wilcoxon rank sum test statistic for site position bias within reads was at least -8, and the read depth over all samples was not larger than 1,380 (twice the mean read depth). For the mitochondrial scaffold, the maximum depth filter was excluded. Individual genotypes with a read depth below 10 or a genotype quality below 20 were set to missing. Monomorphic sites, multiallelic sites, single-nucleotide polymorphisms (SNPs) within 5 bp of indels, and indels themselves were removed. Finally, a two-sided binomial test on allelic balance was performed to ensure high confidence in heterozygous calls, under the null hypothesis that the alternative allele frequency is 0.5. SNPs with *p*-values less than 0.05 were excluded. From 44,072,404 called autosomal variants, 27,530,833 passed hard quality filtering. After applying minimum depth and genotype quality thresholds, and removing indels, monomorphic, and multiallelic sites 3,251,519 SNPs remained. Filtering for a maximum of 10% missing data reduced this to 2,842,750 SNPs. Applying the allelic balance filter yielded a final set of 2,690,649 SNPs. Genome-wide variation was assessed by performing principal component analyses (PCA) using the smartpca function in EIGENSOFT v.7.2.1 (Patterson et al., 2006). Variant statistics were generated with BCFtools stats.

### Genetic relatedness

Genetic relatedness based on proportions of inherited alleles within families (identity-by-descent, IBD) was assessed using PLINK v1.9.0-b.7.6 (Purcell et al., 2007) with the option - -genome rel-check. Prior to this, the 2,690,649 SNP-set was linkage disequilibrium (LD) pruned to obtain independent markers (5 kb window, 10 SNP step, *r*^2^ > 0.8), removing 2,083,602 variants in LD.

We used the same linkage-pruned dataset, ensuring violations were assessed at statistically independent loci, to calculate the proportion of shared genotypes within families (*M*_xy_; Blouin et al., 1996) and to identify Mendelian violations. Genotypes were formatted using PLINK (--recode A) so that alleles were coded as 0 = homozygous reference, 1 = heterozygous, and 2 = homozygous alternative. We then computed *M*_xy_ as the number of identical genotypes divided by the total number of valid genotypes compared. Mendelian violations were assessed by comparing offspring genotypes with those permitted under Mendelian inheritance given the available parental genotypes. For uniparental calculations, offspring were required to carry only the maternal allele at sites where the mother is homozygous (0/0 or 2/2); any genotype containing a non-maternal allele was scored as a violation. When both parents were genotyped, offspring genotypes were evaluated against the set of Mendelian-compatible genotypes for each parental genotype combination, with any incompatible genotype classified as a violation.

### Genome-wide and chromosome-wide heterozygosity retention

To obtain an understanding of the reproductive mode of the uniparental *Cy. frontosa* family, the genome-wide retained heterozygosity was assessed using a custom Bash script. In short, the script operates as follows, using the 2,690,649 SNP-set as input: (i) Heterozygous sites were selected in the parent(s). For the uniparental family, only variant sites where the mother was heterozygous were selected, whereas for the biparental families, only sites where at least one parent was heterozygous were selected. This automatically excluded missing sites in the uniparental family but included them in the biparental families if the one parent was heterozygous (ii) For each offspring, the selected sites were inspected for missing genotypes. If a site was missing in an offspring, it was excluded from that offspring’s denominator (informative sites). (iii) After generating the denominator, three numerators were calculated by counting sites where the offspring’s genotype was heterozygous or homozygous reference or alternative (iv) Percentage values on heterozygosity retention/loss were reported. The chromosome-wide heterozygosity retention analysis aimed to quantify and visualize the distribution of retained/lost heterozygous sites along each chromosome. For that, a custom Bash script similar to the script above was applied. The two main differences were as follows: (i) chromosome VCFs were analyzed instead of the concatenated VCF, and (ii) every informative position and its genotype class, together with the corresponding position was recorded. The distribution of heterozygosity retention and loss along each chromosome was visualized using a custom R-script in the R Statistical Software (v4.5.1; R Core Team, 2025).

### Runs of homozygosity

Runs of homozygosity (ROH) were analyzed per chromosome for each individual using the R package detectRUNS (v0.9.6; Biscarini et al., 2019) to identify and visualize long homozygous tracts and to compare homozygosity patterns among families across chromosomes. Sliding-window-based detection was performed with the following parameters: windowSize = 10, threshold = 0.05, minSNP = 10, ROHet = FALSE, maxOppWindow = 1, maxMissWindow = 1, maxGap = 1e6, minLengthBps = 50,000, minDensity = 1/1e3, maxOppRun = NULL, maxMissRun = NULL. The same parameters were applied to all chromosomes for consistency, regardless of chromosome length and SNP density. The results were plotted for each chromosome, and centromere positions estimated in Böhne et al. (2023) were added where available.

### Heterozygosity retention around centromeric regions

To quantify heterozygosity retention in centromeric regions, the output of the chromosome-wide heterozygosity retention analysis was used together with information on chromosome lengths and centromere positions (Böhne et al., 2023). Chromosomes without an estimated centromere position were excluded.

Using a custom R script, three different maximum half-window sizes (0.5, 1, and 1.5 Mb) were applied. The script selects the minimum of the following three values as the effective half-window size, according to the position of the centromere: (i) the specified maximum half-window size, (ii) the distance from the centromere to the chromosome start, and (iii) the distance from the centromere to the chromosome end. It then calculates the retained heterozygosity rate within the centromeric region and generates a heatmap to visualize these rates.

## Supplementary Material

Supplementary material accompanies this manuscript.

## Acknowledgements

We thank the late G. Kazumbae and team at Lake Tanganyika for help during fieldwork and F. Ronco for help during sampling; C. Beisel and team at the Genomics Facility Basel at the ETH Zurich Department of Biosystems Science and Engineering (D-BSSE), Basel, for assistance with next-generation sequencing; J. Johnson for fish illustrations in Fig. 2. Calculations were performed at sciCORE (http://scicore.unibas.ch/) scientific computing center at the University of Basel. AI-assisted tools were used for language editing (proofreading) and code debugging. All outputs were reviewed and validated by the authors.

## Author contributions

Conceptualization: JMIB, WS; Data curation: JMIB, DSC, MEU; Formal analysis: JMIB, MEU; Funding acquisition: JMIB, WS; Investigation: JMIB, MEU; Project administration: JMIB; Resources: AM, AI, WG, WS; Supervision: JMIB, WS; Validation: JMIB; Visualization: JMIB, MEU; Writing – original draft: MEU, JMIB; All authors have contributed to writing – review & editing.

## Funding

This research was funded the European Union’s Horizon 2020 research and innovation programme under the Marie Skłodowska-Curie grant agreement 891088 to JMIB, and by the Swiss National Science Foundation (SNSF) grant 189897 within the COST (European Cooperation in Science and Technology) funding scheme to JMIB and WS.

## Data availability

Raw sequence data are deposited on the NCBI database under BioProject: PRJNA798382. Code for computational analyses is available from Github: https://github.com/JMIBarth/cfrontosa.

## Competing interests

The authors declare no competing interests.

## Notes

### Competing Interest Statement

The authors have declared no competing interest.

