## Supplementary Material for "Genomic evidence for facultative selfing in the cichlid fish *Cyphotilapia frontosa*"

**Supplementary table 1: Per-sample variant metrics derived from biallelic SNPs.** Counts (n) of homozygous reference (ref hom), alternative homozygous (alt hom), and heterozygous (het) variants. Transitions and transversions, the average depth at variant sites, singletons, and missing variants.

| Fam | ID | Ref hom (n) | Alt hom (n) | Het (n) | Het/alt hom | Het % | Transitions (n) | Transv (n) | Ts/Tv | Avg depth | Singletons (n) | Missing (n) |
|---|---|---|---|---|---|---|---|---|---|---|---|---|
| *Cy. frontosa* (wild) | **Cphfro_M_04G6** | 1382679 | 680577 | 585584 | 0.86 | 22.11 | 882557 | 383604 | 2.30 | 35.9 | 2433 | 41809 |
| | **Cphfro_F_04G7** | 1443726 | 737360 | 481281 | 0.65 | 18.08 | 848989 | 369652 | 2.30 | 37.1 | 416 | 28282 |
| | **Cphfro_1_04G8** | 1472476 | 766301 | 425178 | 0.55 | 15.96 | 829447 | 362032 | 2.29 | 36.1 | 329 | 26694 |
| | **Cphfro_1_04H1** | 1504839 | 798142 | 365042 | 0.46 | 13.68 | 809156 | 354028 | 2.29 | 40.8 | 270 | 22626 |
| | **Cphfro_1_04H2** | 1463607 | 755983 | 444803 | 0.59 | 16.69 | 836362 | 364424 | 2.30 | 40.5 | 493 | 26256 |
| | **Cphfro_1_04H4** | 1513853 | 807545 | 347267 | 0.43 | 13.01 | 803458 | 351354 | 2.29 | 39.8 | 508 | 21984 |
| | **Cphfro_1_04H5** | 1486104 | 779250 | 401784 | 0.52 | 15.06 | 821903 | 359131 | 2.29 | 39.6 | 347 | 23511 |
| *Ct. benthicola* | **Cteben_M_04F8** | 1531153 | 975576 | 153119 | 0.16 | 5.76 | 782713 | 345982 | 2.26 | 33.9 | 3340 | 30801 |
| | **Cteben_F_04F9** | 1537386 | 979612 | 150599 | 0.15 | 5.65 | 783651 | 346560 | 2.26 | 32.9 | 2135 | 23052 |
| | **Cteben_1_04G1** | 1540361 | 979556 | 153525 | 0.16 | 5.74 | 785487 | 347594 | 2.26 | 35.0 | 220 | 17207 |
| | **Cteben_1_04G2** | 1544136 | 983467 | 147206 | 0.15 | 5.50 | 783817 | 346856 | 2.26 | 39.1 | 269 | 15840 |
| | **Cteben_1_04G3** | 1539825 | 978690 | 154828 | 0.16 | 5.79 | 785875 | 347643 | 2.26 | 35.1 | 327 | 17306 |
| | **Cteben_1_04G4** | 1539343 | 980949 | 149117 | 0.15 | 5.59 | 783514 | 346552 | 2.26 | 32.6 | 335 | 21240 |
| | **Cteben_1_04G5** | 1528776 | 974504 | 155763 | 0.16 | 5.86 | 783689 | 346578 | 2.26 | 30.7 | 417 | 31606 |
| *Cy. frontosa* (uniparental) | **Cphfro_F_AM01** | 1478512 | 777172 | 411284 | 0.53 | 15.42 | 827976 | 360480 | 2.30 | 65.2 | 132 | 23681 |
| | **Cphfro_1_AM03** | 1405593 | 798429 | 167390 | 0.21 | 7.06 | 675282 | 290537 | 2.32 | 30.9 | 377 | 319237 |
| | **Cphfro_1_AM04** | 1563725 | 869483 | 231776 | 0.27 | 8.70 | 766186 | 335073 | 2.29 | 33.0 | 107 | 25665 |
| | **Cphfro_1_AM05** | 1581026 | 886916 | 197441 | 0.22 | 7.41 | 753838 | 330519 | 2.28 | 36.7 | 120 | 25266 |
| | **Cphfro_1_AM06** | 1572150 | 875335 | 222901 | 0.25 | 8.35 | 763342 | 334894 | 2.28 | 34.3 | 195 | 20263 |
| | **Cphfro_1_AM07** | 1565255 | 866878 | 241773 | 0.28 | 9.04 | 770953 | 337698 | 2.28 | 40.2 | 59 | 16743 |
| | **Cphfro_1_AM08** | 1592643 | 900597 | 168502 | 0.19 | 6.33 | 742718 | 326381 | 2.28 | 33.5 | 148 | 28907 |

**Supplementary table 2: Identity-by-descent (IBD).** Pairwise (ID1 and ID2) within-family comparisons of IBD. Relationship types (RT) were inferred by PLINK as parent-offspring (PO), full sibling (FS), or other (OT) from the probability that individual pairs share zero (Z0), one (Z1), or two alleles (Z2) IBD, respectively, the proportion of the genome shared (PI_HAT), and the excess IBD Z-score (EZ).

| Fam | ID1 | ID2 | RT | EZ | Z0 | Z1 | Z2 | PI_ HAT |
|---|---|---|---|---|---|---|---|---|
| *Cy. frontosa* (wild) | **1_04G8** | **1_04H1** | FS | 0.5 | 0.1244 | 0.4128 | 0.4628 | 0.6692 |
| | **1_04G8** | **1_04H2** | FS | 0.5 | 0.0668 | 0.4162 | 0.5171 | 0.7251 |
| | **1_04G8** | **1_04H4** | FS | 0.5 | 0.1940 | 0.3951 | 0.4109 | 0.6084 |
| | **1_04G8** | **1_04H5** | FS | 0.5 | 0.1368 | 0.3418 | 0.5214 | 0.6923 |
| | **1_04G8** | **F_04G7** | PO | 0.5 | 0.0047 | 0.4586 | 0.5367 | 0.7660 |
| | **1_04G8** | **M_04G6** | PO | 0.5 | 0.0058 | 0.5787 | 0.4155 | 0.7048 |
| | **1_04H1** | **1_04H2** | FS | 0.5 | 0.1257 | 0.4769 | 0.3974 | 0.6358 |
| | **1_04H1** | **1_04H4** | FS | 0.5 | 0.2273 | 0.2698 | 0.5029 | 0.6378 |
| | **1_04H1** | **1_04H5** | FS | 0.5 | 0.1514 | 0.3650 | 0.4836 | 0.6661 |
| | **1_04H1** | **F_04G7** | PO | 0.5 | 0.0058 | 0.5084 | 0.4858 | 0.7400 |
| | **1_04H1** | **M_04G6** | PO | 0.5 | 0.0047 | 0.5887 | 0.4065 | 0.7009 |
| | **1_04H2** | **1_04H4** | FS | 0.5 | 0.1537 | 0.3387 | 0.5076 | 0.6770 |
| | **1_04H2** | **1_04H5** | FS | 0.5 | 0.1244 | 0.4875 | 0.3882 | 0.6319 |
| | **1_04H2** | **F_04G7** | PO | 0.5 | 0.0052 | 0.5789 | 0.4160 | 0.7054 |
| | **1_04H2** | **M_04G6** | PO | 0.5 | 0.0055 | 0.4679 | 0.5266 | 0.7606 |
| | **1_04H4** | **1_04H5** | FS | 0.5 | 0.3279 | 0.1842 | 0.4879 | 0.5800 |
| | **1_04H4** | **F_04G7** | PO | 0.5 | 0.0047 | 0.5614 | 0.4338 | 0.7146 |
| | **1_04H4** | **M_04G6** | PO | 0.5 | 0.0053 | 0.5571 | 0.4375 | 0.7161 |
| | **1_04H5** | **F_04G7** | PO | 0.5 | 0.0036 | 0.3514 | 0.6450 | 0.8207 |
| | **1_04H5** | **M_04G6** | PO | 0.5 | 0.0067 | 0.7243 | 0.2690 | 0.6312 |
| *Cy. frontosa* (uniparental) | **1_AM03** | **1_AM04** | FS | 0.25 | 0.2089 | 0.0000 | 0.7911 | 0.7911 |
| | **1_AM03** | **1_AM05** | FS | 0.25 | 0.1893 | 0.0000 | 0.8107 | 0.8107 |
| | **1_AM03** | **1_AM06** | FS | 0.25 | 0.1829 | 0.0618 | 0.7553 | 0.7862 |
| | **1_AM03** | **1_AM07** | FS | 0.25 | 0.0904 | 0.1204 | 0.7892 | 0.8494 |
| | **1_AM03** | **1_AM08** | FS | 0.25 | 0.2675 | 0.0000 | 0.7325 | 0.7325 |
| | **1_AM03** | **F_AM01** | OT | 0 | 0.0027 | 0.2332 | 0.7641 | 0.8807 |
| | **1_AM04** | **1_AM05** | FS | 0.25 | 0.1742 | 0.0000 | 0.8258 | 0.8258 |
| | **1_AM04** | **1_AM06** | FS | 0.25 | 0.2509 | 0.0000 | 0.7491 | 0.7491 |
| | **1_AM04** | **1_AM07** | FS | 0.25 | 0.1475 | 0.0141 | 0.8384 | 0.8454 |
| | **1_AM04** | **1_AM08** | FS | 0.25 | 0.2158 | 0.0000 | 0.7842 | 0.7842 |
| | **1_AM04** | **F_AM01** | OT | 0 | 0.0034 | 0.1868 | 0.8098 | 0.9032 |
| | **1_AM05** | **1_AM06** | FS | 0.25 | 0.2308 | 0.0021 | 0.7671 | 0.7681 |

| Fam | ID1 | ID2 | RT | EZ | Z0 | Z1 | Z2 | PI_ HAT |
|---|---|---|---|---|---|---|---|---|
| *Cy. frontosa* (uniparental) | **1_AM05** | **1_AM07** | FS | 0.25 | 0.0281 | 0.2383 | 0.7336 | 0.8528 |
| | **1_AM05** | **1_AM08** | FS | 0.25 | 0.2677 | 0.0000 | 0.7323 | 0.7323 |
| | **1_AM05** | **F_AM01** | OT | 0 | 0.0027 | 0.2130 | 0.7843 | 0.8908 |
| | **1_AM06** | **1_AM07** | FS | 0.25 | 0.2212 | 0.0000 | 0.7788 | 0.7788 |
| | **1_AM06** | **1_AM08** | FS | 0.25 | 0.3310 | 0.0000 | 0.6690 | 0.6690 |
| | **1_AM06** | **F_AM01** | OT | 0 | 0.0030 | 0.2108 | 0.7861 | 0.8915 |
| | **1_AM07** | **1_AM08** | FS | 0.25 | 0.3718 | 0.0000 | 0.6282 | 0.6282 |
| | **1_AM07** | **F_AM01** | OT | 0 | 0.0026 | 0.1838 | 0.8136 | 0.9055 |
| | **1_AM08** | **F_AM01** | OT | 0 | 0.0028 | 0.2551 | 0.7421 | 0.8696 |
| | **F_04G7** | **M_04G6** | OT | 0 | 0.0087 | 0.8733 | 0.1180 | 0.5546 |
| *Ct. benthicola* | **1_04G1** | **1_04G2** | FS | 0.5 | 0.0081 | 0.0991 | 0.8928 | 0.9423 |
| | **1_04G1** | **1_04G3** | FS | 0.5 | 0.0074 | 0.1080 | 0.8846 | 0.9386 |
| | **1_04G1** | **1_04G4** | FS | 0.5 | 0.0072 | 0.1242 | 0.8686 | 0.9307 |
| | **1_04G1** | **1_04G5** | FS | 0.5 | 0.0133 | 0.1200 | 0.8667 | 0.9267 |
| | **1_04G1** | **F_04F9** | PO | 0.5 | 0.0040 | 0.1863 | 0.8097 | 0.9029 |
| | **1_04G1** | **M_04F8** | PO | 0.5 | 0.0042 | 0.1867 | 0.8091 | 0.9025 |
| | **1_04G2** | **1_04G3** | FS | 0.5 | 0.0074 | 0.1251 | 0.8676 | 0.9301 |
| | **1_04G2** | **1_04G4** | FS | 0.5 | 0.0084 | 0.1163 | 0.8753 | 0.9334 |
| | **1_04G2** | **1_04G5** | FS | 0.5 | 0.0170 | 0.1297 | 0.8534 | 0.9182 |
| | **1_04G2** | **F_04F9** | PO | 0.5 | 0.0040 | 0.1809 | 0.8151 | 0.9056 |
| | **1_04G2** | **M_04F8** | PO | 0.5 | 0.0042 | 0.1929 | 0.8029 | 0.8993 |
| | **1_04G3** | **1_04G4** | FS | 0.5 | 0.0077 | 0.1116 | 0.8807 | 0.9365 |
| | **1_04G3** | **1_04G5** | FS | 0.5 | 0.0134 | 0.1197 | 0.8669 | 0.9267 |
| | **1_04G3** | **F_04F9** | PO | 0.5 | 0.0037 | 0.1845 | 0.8117 | 0.9040 |
| | **1_04G3** | **M_04F8** | PO | 0.5 | 0.0041 | 0.1885 | 0.8074 | 0.9016 |
| | **1_04G4** | **1_04G5** | FS | 0.5 | 0.0114 | 0.1091 | 0.8795 | 0.9341 |
| | **1_04G4** | **F_04F9** | PO | 0.5 | 0.0039 | 0.1867 | 0.8094 | 0.9028 |
| | **1_04G4** | **M_04F8** | PO | 0.5 | 0.0035 | 0.1857 | 0.8107 | 0.9036 |
| | **1_04G5** | **F_04F9** | PO | 0.5 | 0.0032 | 0.1833 | 0.8135 | 0.9051 |
| | **1_04G5** | **M_04F8** | PO | 0.5 | 0.0040 | 0.1937 | 0.8022 | 0.8991 |
| | **F_04F9** | **M_04F8** | OT | 0 | 0.2582 | 0.0000 | 0.7418 | 0.7418 |

**Supplementary table 3: Pairwise (ID1 and ID2) within-family comparisons of genotype similarity ($M_{xy}$).** $M_{xy}$ was calculated as the number of identical genotypes (n_loci_identical) divided by the total number of valid genotypes compared (n_loci_compared). Missing data was excluded.

| Fam | ID1 | ID2 | Mxy | Loci identical (n) | Loci compared (n) |
|---|---|---|---|---|---|
| *Cy. frontosa* (wild) | 1_04G8 | 1_04H1 | 0.76664 | 453925 | 592095 |
| | 1_04G8 | 1_04H2 | 0.79742 | 471148 | 590842 |
| | 1_04G8 | 1_04H4 | 0.73492 | 435266 | 592262 |
| | 1_04G8 | 1_04H5 | 0.78799 | 466275 | 591729 |
| | 1_04H1 | 1_04H2 | 0.74046 | 438186 | 591773 |
| | 1_04H1 | 1_04H4 | 0.76621 | 454649 | 593376 |
| | 1_04H1 | 1_04H5 | 0.77063 | 456790 | 592748 |
| | 1_04H2 | 1_04H4 | 0.77982 | 461701 | 592062 |
| | 1_04H2 | 1_04H5 | 0.73702 | 435861 | 591384 |
| | 1_04H4 | 1_04H5 | 0.74419 | 441195 | 592849 |
| | F_04G7 | M_04G6 | 0.64807 | 383291 | 591435 |
| | 1_04G8 | F_04G7 | 0.81512 | 483802 | 593533 |
| | 1_04G8 | M_04G6 | 0.76678 | 451727 | 589124 |
| | 1_04H1 | F_04G7 | 0.79471 | 472481 | 594533 |
| | 1_04H1 | M_04G6 | 0.76338 | 450514 | 590155 |
| | 1_04H2 | F_04G7 | 0.76707 | 455028 | 593206 |
| | 1_04H2 | M_04G6 | 0.81098 | 477535 | 588836 |
| | 1_04H4 | F_04G7 | 0.77424 | 460453 | 594714 |
| | 1_04H4 | M_04G6 | 0.77561 | 457876 | 590344 |
| | 1_04H5 | F_04G7 | 0.85835 | 510083 | 594258 |
| | 1_04H5 | M_04G6 | 0.70842 | 417735 | 589675 |
| *Cy. frontosa* (uniparental) | 1_AM03 | 1_AM04 | 0.90355 | 473264 | 523785 |
| | 1_AM03 | 1_AM05 | 0.92113 | 482921 | 524268 |
| | 1_AM03 | 1_AM06 | 0.87360 | 458029 | 524302 |
| | 1_AM03 | 1_AM07 | 0.90182 | 473470 | 525017 |
| | 1_AM03 | 1_AM08 | 0.89278 | 467658 | 523820 |
| | 1_AM04 | 1_AM05 | 0.91160 | 541312 | 593805 |
| | 1_AM04 | 1_AM06 | 0.88278 | 525561 | 595350 |
| | 1_AM04 | 1_AM07 | 0.91224 | 543965 | 596299 |
| | 1_AM04 | 1_AM08 | 0.90241 | 534883 | 592725 |
| | 1_AM05 | 1_AM06 | 0.87062 | 518330 | 595359 |
| | 1_AM05 | 1_AM07 | 0.88966 | 530495 | 596291 |
| *Cy. frontosa* (uniparental) | 1_AM05 | 1_AM08 | 0.90022 | 533609 | 592753 |
| | 1_AM06 | 1_AM07 | 0.90574 | 541542 | 597898 |
| | 1_AM06 | 1_AM08 | 0.88611 | 526584 | 594263 |
| | 1_AM07 | 1_AM08 | 0.87576 | 521232 | 595178 |
| | 1_AM03 | F_AM01 | 0.90582 | 475815 | 525289 |
| | 1_AM04 | F_AM01 | 0.92388 | 551339 | 596764 |
| | 1_AM05 | F_AM01 | 0.91386 | 545436 | 596847 |
| | 1_AM06 | F_AM01 | 0.91451 | 547226 | 598379 |
| | 1_AM07 | F_AM01 | 0.92549 | 554690 | 599350 |
| | 1_AM08 | F_AM01 | 0.89705 | 534356 | 595684 |
| *Ct. benthicola* | 1_04G1 | 1_04G2 | 0.95609 | 571278 | 597517 |
| | 1_04G1 | 1_04G3 | 0.95297 | 569075 | 597159 |
| | 1_04G1 | 1_04G4 | 0.94664 | 564324 | 596131 |
| | 1_04G1 | 1_04G5 | 0.94490 | 560691 | 593384 |
| | 1_04G2 | 1_04G3 | 0.94619 | 565301 | 597449 |
| | 1_04G2 | 1_04G4 | 0.94910 | 566044 | 596400 |
| | 1_04G2 | 1_04G5 | 0.93901 | 557455 | 593661 |
| | 1_04G3 | 1_04G4 | 0.95134 | 567076 | 596081 |
| | 1_04G3 | 1_04G5 | 0.94496 | 560661 | 593315 |
| | 1_04G4 | 1_04G5 | 0.95030 | 562923 | 592364 |
| | F_04F9 | M_04F8 | 0.86933 | 514824 | 592206 |
| | 1_04G1 | F_04F9 | 0.92376 | 550498 | 595934 |
| | 1_04G1 | M_04F8 | 0.92346 | 548169 | 593602 |
| | 1_04G2 | F_04F9 | 0.92588 | 551964 | 596148 |
| | 1_04G2 | M_04F8 | 0.92099 | 546966 | 593889 |
| | 1_04G3 | F_04F9 | 0.92457 | 550871 | 595810 |
| | 1_04G3 | M_04F8 | 0.92280 | 547706 | 593529 |
| | 1_04G4 | F_04F9 | 0.92365 | 549409 | 594826 |
| | 1_04G4 | M_04F8 | 0.92421 | 547670 | 592580 |
| | 1_04G5 | F_04F9 | 0.92536 | 547992 | 592193 |
| | 1_04G5 | M_04F8 | 0.92076 | 543111 | 589849 |

**Supplementary table 4: Mendelian violations.** Mendelian violations assessed by two methods: uniparental test (offspring vs. mother) and trio test (offspring vs. both parents) for both, the linkage-pruned and full variant datasets.

| | | | Linkage pruned (607'047 SNPs) | | | | | |
|---|---|---|---|---|---|---|---|---|
| **Fam** | **ID Female** | **ID Male** | **Offspring (n)** | **Total loci** | **Violations female_only** | **Violation rate (female_only)** | **Violations trio** | **Violation rate trio** |
| ***Cy. frontosa* (wild)** | **Cphfro_F_04G7** | **Cphfro_M_04G6** | 5 | 2'970'244 | 274'031 | 0.09226 | 2'359 | 0.00079 |
| ***Ct. benthicola*** | **Cteben_F_04F9** | **Cteben_M_04F8** | 5 | 2'974'911 | 136'616 | 0.04592 | 1'947 | 0.00065 |
| ***Cy. frontosa* (uniparental)** | **Cphfro_F_AM01** | **NA** | 6 | 3'512'313 | 1'573 | 0.00045 | NA | NA |
| | | | **Full dataset (2,690,649 SNPs)** | | | | | |
| ***Cy. frontosa* (wild)** | **Cphfro_F_04G7** | **Cphfro_M_04G6** | 5 | 13'194'813 | 709'870 | 0.05380 | 4'640 | 0.00035 |
| ***Ct. benthicola*** | **Cteben_F_04F9** | **Cteben_M_04F8** | 5 | 13'239'238 | 355'505 | 0.02685 | 4'022 | 0.00030 |
| ***Cy. frontosa* (uniparental)** | **Cphfro_F_AM01** | **NA** | 6 | 15'571'754 | 4'111 | 0.00026 | NA | NA |

**Supplementary table 5: Genome-wide heterozygosity retention.** Counts (n) of informative sites in the offspring where the mother is heterozygous in the uniparental family and at least one parent is heterozygous in the biparental families (inform), sites homozygous reference (ref hom), sites alternative homozygous (alt hom), and retained heterozygous sites (het retained), as well as their percentages.

| Fam | ID offspring | Inform (n) | Ref hom (n) | Ref hom % | Alt hom (n) | Alt hom % | Het retained (n) | Het retained % |
|---|---|---|---|---|---|---|---|---|
| ***Cy. frontosa* (wild)** | **Cphfro_1_04G8** | 784'375 | 210'923.00 | 26.89 | 160'913 | 20.51 | 412'539 | 52.59 |
| | **Cphfro_1_04H1** | 786'328 | 240'585.00 | 30.60 | 191'612 | 24.37 | 354'131 | 45.04 |
| | **Cphfro_1_04H2** | 782'855 | 201'023.00 | 25.68 | 151'125 | 19.30 | 430'707 | 55.02 |
| | **Cphfro_1_04H4** | 787'260 | 249'908.00 | 31.74 | 201'280 | 25.57 | 336'072 | 42.69 |
| | **Cphfro_1_04H5** | 785'658 | 222'874.00 | 28.37 | 173'256 | 22.05 | 389'528 | 49.58 |
| ***Ct. benthicola*** | **Cteben_1_04G1** | 247'223 | 67'723.00 | 27.39 | 54'533 | 22.06 | 124'967 | 50.55 |
| | **Cteben_1_04G2** | 247'201 | 70'416.00 | 28.49 | 57'950 | 23.44 | 118'835 | 48.07 |
| | **Cteben_1_04G3** | 246'805 | 67'098.00 | 27.19 | 53'549 | 21.70 | 126'158 | 51.12 |
| | **Cteben_1_04G4** | 246'567 | 69'323.00 | 28.12 | 56'467 | 22.90 | 120'777 | 48.98 |
| | **Cteben_1_04G5** | 245'997 | 66'213.00 | 26.92 | 53'002 | 21.55 | 126'782 | 51.54 |
| ***Cy. frontosa* (uniparental)** | **Cphfro_1_AM03** | 356'674 | 98'596.00 | 27.64 | 100'229 | 28.10 | 157'849 | 44.26 |
| | **Cphfro_1_AM04** | 400'226 | 90'013.00 | 22.49 | 90'232 | 22.55 | 219'981 | 54.96 |
| | **Cphfro_1_AM05** | 400'759 | 106'711.00 | 26.63 | 107'211 | 26.75 | 186'837 | 46.62 |
| | **Cphfro_1_AM06** | 400'593 | 94'488.00 | 23.59 | 94'800 | 23.66 | 211'305 | 52.75 |
| | **Cphfro_1_AM07** | 400'478 | 85'192.00 | 21.27 | 85'791 | 21.42 | 229'495 | 57.31 |
| | **Cphfro_1_AM08** | 401'589 | 120'738.00 | 30.07 | 121'503 | 30.26 | 159'348 | 39.68 |

**Supplementary table 6: Centromere positions.** Chromosome (linkage group; LG) and RefSeq (GCF_001858045.2) IDs, chromosome lengths (bp), and centromere positions from Böhne et al., 2023.

| RefSeq | Chromosome | Size (bp) | Centromere position |
|---|---|---|---|
| **NC_031965.2** | **LG1** | 40673430 | NA |
| **NC_031966.2** | **LG2** | 36523203 | NA |
| **NC_031967.2** | **LG3** | 87567345 | 3616177 |
| **NC_031969.2** | **LG4** | 35549522 | 2639573 |
| **NC_031970.2** | **LG5** | 39714817 | 38013903 |
| **NC_031971.2** | **LG6** | 42433576 | 949777 |
| **NC_031972.2** | **LG7** | 64772279 | 8582623 |
| **NC_031973.2** | **LG8** | 30527416 | 29369617 |
| **NC_031974.2** | **LG9** | 35850837 | 30300790.75 |
| **NC_031975.2** | **LG10** | 34704454 | NA |
| **NC_031976.2** | **LG11** | 39275952 | 4179490 |
| **NC_031977.2** | **LG12** | 38600464 | NA |
| **NC_031978.2** | **LG13** | 34734273 | 32424445 |
| **NC_031979.2** | **LG14** | 40509636 | 4927716 |
| **NC_031980.2** | **LG15** | 39688505 | 31537495 |
| **NC_031981.2** | **LG17** | 38839487 | 35654356 |
| **NC_031982.2** | **LG18** | 38636442 | 35950441 |
| **NC_031983.2** | **LG19** | 30963196 | 525320.5 |
| **NC_031984.2** | **LG20** | 37140374 | NA |
| **NC_031985.2** | **LG22** | 39199643 | NA |

**Supplementary table 7: Heterozygosity retention.** Counts (n) of SNPs and retained heterozygous sites (%) within centromere-centered windows per chromosome and per offspring of the uniparental *Cy. frontosa* family for three maximum half-window sizes of 1, 0.5, and 1.5 Mb. Chromosomes without an estimated centromere position were excluded.

| | **Maximum half-window size: 1 Mb** | | | | | | | | | | | |
|---|---|---|---|---|---|---|---|---|---|---|---|---|
| **ID** | **AM03** | | **AM04** | | **AM05** | | **AM06** | | **AM07** | | **AM08** | |
| **Chromosome** | **SNPs (n)** | **Het (%)** | **SNPs (n)** | **Het (%)** | **SNPs (n)** | **Het (%)** | **SNPs (n)** | **Het (%)** | **SNPs (n)** | **Het (%)** | **SNPs (n)** | **Het (%)** |
| **LG3** | 620 | 100.0 | 726 | 1.4 | 721 | 1.1 | 717 | 1.3 | 727 | 1.0 | 723 | 1.4 |
| **LG4** | 229 | 9.2 | 240 | 10.8 | 239 | 10.0 | 237 | 9.3 | 231 | 99.6 | 239 | 9.6 |
| **LG5** | 683 | 100.0 | 751 | 100.0 | 745 | 100.0 | 777 | 0.3 | 754 | 100.0 | 775 | 0.3 |
| **LG6** | 25 | 76.0 | 50 | 58.0 | 47 | 95.7 | 45 | 77.8 | 49 | 57.1 | 51 | 76.5 |
| **LG7** | 458 | 1.1 | 514 | 99.8 | 527 | 0.9 | 526 | 0.6 | 531 | 0.9 | 522 | 0.6 |
| **LG8** | 412 | 99.8 | 463 | 2.6 | 447 | 100.0 | 463 | 1.9 | 451 | 100.0 | 450 | 70.0 |
| **LG9** | 364 | 1.1 | 396 | 99.7 | 414 | 1.0 | 402 | 99.8 | 416 | 1.0 | 393 | 99.7 |
| **LG11** | 287 | 100.0 | 344 | 4.9 | 345 | 4.9 | 341 | 4.7 | 342 | 4.1 | 343 | 5.2 |
| **LG13** | 365 | 99.2 | 414 | 1.0 | 406 | 100.0 | 417 | 0.5 | 411 | 0.7 | 398 | 100.0 |
| **LG14** | 172 | 97.7 | 200 | 98.0 | 193 | 98.4 | 200 | 47.0 | 193 | 99.5 | 194 | 99.5 |
| **LG15** | 162 | 13.0 | 171 | 100.0 | 162 | 100.0 | 173 | 11.6 | 173 | 100.0 | 174 | 12.6 |
| **LG17** | 399 | 0.8 | 449 | 0.2 | 437 | 100.0 | 454 | 0.7 | 451 | 14.9 | 444 | 99.8 |
| **LG18** | 360 | 100.0 | 409 | 52.1 | 406 | 100.0 | 406 | 3.0 | 404 | 100.0 | 404 | 3.0 |
| **LG19** | 136 | 6.6 | 139 | 6.5 | 140 | 6.4 | 130 | 100.0 | 136 | 100.0 | 140 | 6.4 |
| **LG23** | 229 | 98.3 | 256 | 96.9 | 254 | 98.4 | 262 | 3.4 | 255 | 37.3 | 261 | 2.3 |
| **LG16** | 18 | 61.1 | 19 | 63.2 | 16 | 81.2 | 19 | 84.2 | 19 | 63.2 | 15 | 100.0 |
| | **Maximum half-window size: 0.5 Mb** | | | | | | | | | | | |
| **LG3** | 300 | 100.0 | 341 | 2.6 | 339 | 2.4 | 337 | 2.4 | 341 | 1.8 | 340 | 2.6 |
| **LG4** | 61 | 8.2 | 63 | 12.7 | 63 | 11.1 | 63 | 12.7 | 59 | 98.3 | 63 | 12.7 |
| **LG5** | 350 | 100.0 | 391 | 100.0 | 398 | 100.0 | 411 | 0.2 | 402 | 100.0 | 412 | 0.2 |
| **LG6** | 14 | 100.0 | 34 | 79.4 | 33 | 97.0 | 32 | 81.2 | 34 | 79.4 | 34 | 79.4 |
| **LG7** | 268 | 1.9 | 292 | 100.0 | 300 | 1.7 | 299 | 1.0 | 303 | 1.7 | 298 | 1.0 |
| **LG8** | 121 | 99.2 | 132 | 7.6 | 129 | 100.0 | 129 | 7.0 | 127 | 100.0 | 125 | 88.0 |
| **LG9** | 110 | 3.6 | 125 | 100.0 | 132 | 3.0 | 131 | 100.0 | 133 | 3.0 | 125 | 100.0 |
| **LG11** | 64 | 100.0 | 89 | 15.7 | 88 | 14.8 | 85 | 16.5 | 85 | 12.9 | 87 | 16.1 |
| **LG13** | 158 | 98.7 | 186 | 1.1 | 186 | 100.0 | 188 | 0.5 | 185 | 1.1 | 181 | 100.0 |
| **LG14** | 42 | 92.9 | 55 | 94.5 | 52 | 94.2 | 54 | 7.4 | 49 | 100.0 | 53 | 100.0 |
| **LG15** | 46 | 30.4 | 47 | 100.0 | 44 | 100.0 | 48 | 27.1 | 47 | 100.0 | 48 | 27.1 |
| **LG17** | 223 | 0.9 | 254 | 0.4 | 244 | 100.0 | 256 | 0.8 | 256 | 2.7 | 249 | 100.0 |
| **LG18** | 163 | 100.0 | 183 | 9.8 | 177 | 100.0 | 174 | 2.3 | 174 | 100.0 | 172 | 2.3 |
| **LG19** | 120 | 7.5 | 123 | 7.3 | 124 | 7.3 | 115 | 100.0 | 120 | 100.0 | 124 | 7.3 |
| **LG23** | 113 | 96.5 | 124 | 93.5 | 121 | 96.7 | 124 | 7.3 | 121 | 36.4 | 124 | 4.8 |
| **LG16** | 6 | 100.0 | 7 | 100.0 | 7 | 57.1 | 7 | 57.1 | 7 | 100.0 | 7 | 100.0 |
| | **Maximum half-window size: 1.5 Mb** | | | | | | | | | | | |
| **LG3** | 890 | 99.9 | 1'062 | 2.1 | 1'056 | 1.8 | 1'049 | 1.7 | 1'065 | 1.9 | 1'060 | 2.0 |
| **LG4** | 351 | 6.0 | 408 | 6.4 | 407 | 5.9 | 404 | 5.4 | 394 | 99.7 | 406 | 5.7 |
| **LG5** | 922 | 99.9 | 1'030 | 99.9 | 1'019 | 100.0 | 1'063 | 0.7 | 1'035 | 99.9 | 1'064 | 1.3 |
| **LG6** | 25 | 76.0 | 50 | 58.0 | 47 | 95.7 | 45 | 77.8 | 49 | 57.1 | 51 | 76.5 |
| **LG7** | 687 | 1.0 | 779 | 99.9 | 798 | 0.6 | 799 | 0.6 | 804 | 0.9 | 793 | 0.8 |
| **LG8** | 505 | 99.8 | 566 | 2.1 | 541 | 100.0 | 566 | 1.6 | 551 | 100.0 | 550 | 57.3 |
| **LG9** | 706 | 2.4 | 741 | 99.7 | 786 | 2.2 | 755 | 99.9 | 785 | 2.0 | 753 | 99.9 |
| **LG11** | 461 | 99.8 | 563 | 3.4 | 562 | 3.0 | 556 | 3.2 | 562 | 2.8 | 560 | 3.6 |
| **LG13** | 481 | 99.2 | 546 | 2.4 | 534 | 99.8 | 546 | 2.0 | 543 | 2.2 | 525 | 99.8 |
| **LG14** | 373 | 98.9 | 461 | 99.1 | 452 | 99.3 | 465 | 55.1 | 457 | 99.8 | 448 | 99.8 |
| **LG15** | 386 | 6.5 | 404 | 100.0 | 390 | 100.0 | 414 | 5.8 | 404 | 100.0 | 417 | 7.0 |
| **LG17** | 684 | 4.7 | 774 | 4.1 | 742 | 100.0 | 780 | 4.5 | 772 | 17.0 | 752 | 99.9 |
| **LG18** | 515 | 100.0 | 571 | 65.7 | 572 | 100.0 | 572 | 2.3 | 568 | 100.0 | 570 | 2.3 |
| **LG19** | 136 | 6.6 | 139 | 6.5 | 140 | 6.4 | 130 | 100.0 | 136 | 100.0 | 140 | 6.4 |
| **LG23** | 229 | 98.3 | 256 | 96.9 | 254 | 98.4 | 262 | 3.4 | 255 | 37.3 | 261 | 2.3 |
| **LG16** | 49 | 24.5 | 48 | 85.4 | 47 | 31.9 | 51 | 33.3 | 50 | 86.0 | 48 | 33.3 |

**Supplementary figure 1: Genome wide principal component analysis (PCA),** based on 2,690,649 SNPs. Specimen IDs are given for individuals with distinct positions.

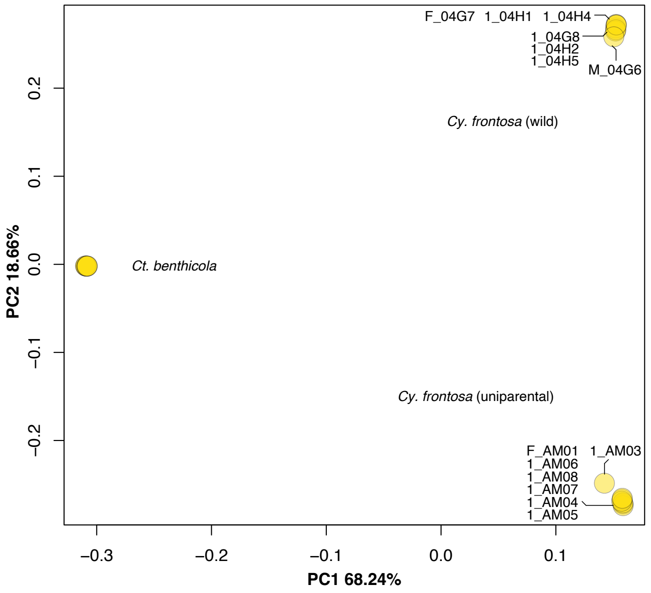


**Supplementary figure 2: Identity-by-descent (IBD) triangle plot** of the probability that individual pairs share one (Z1), or two alleles (Z2) identical by descent. Z1 was plotted instead of Z0 due to the high inbreeding signal and lack of unrelated individuals. Points on the diagonal (Z1+Z2=1) represent where Z0 = 0.

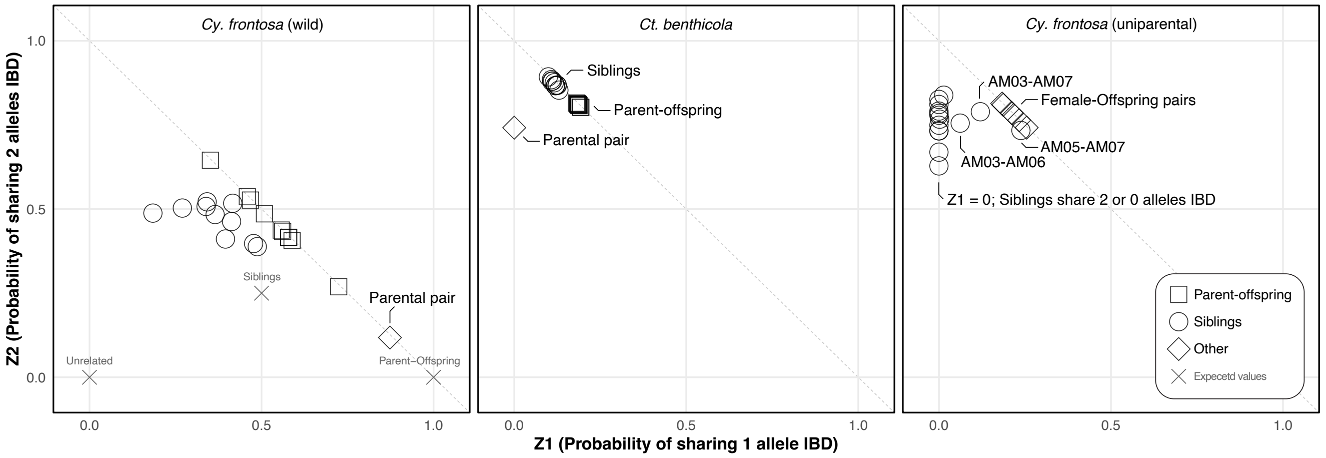

**Supplementary figure 3: Runs of homozygosity (ROH)** across chromosomes per individual for *Cy. frontosa* wild (purple), uniparental (green), and *Ct. benthicola* (blue). Black-filled triangles indicate the estimated centromere positions where available.

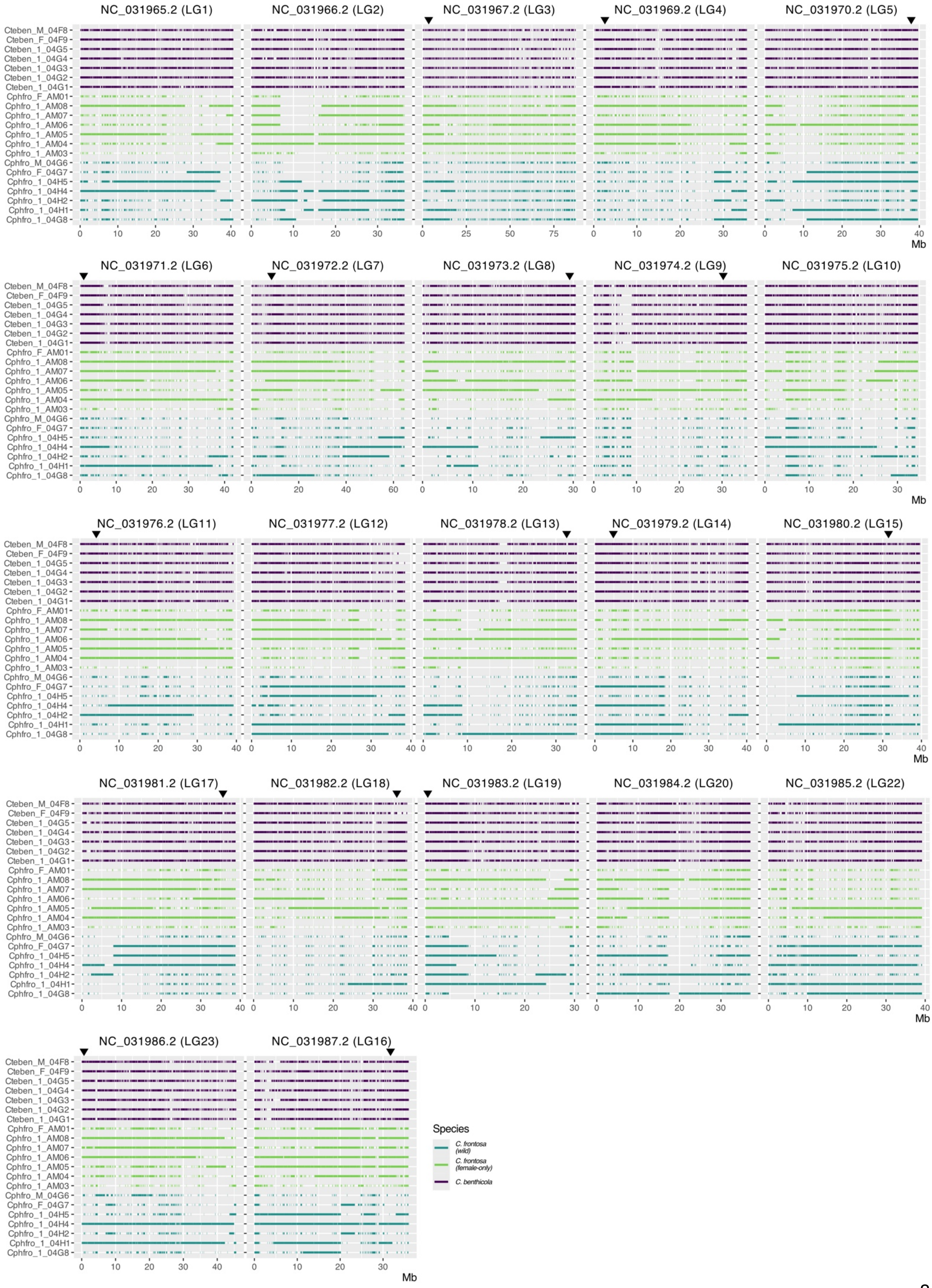

**Supplementary figure 4: Heatmap of heterozygosity retention rates** within centromeric windows across the offspring of the uniparental *Cy. frontosa* family. NC_031987.2 has low SNP density and should be interpreted cautiously. (A) Centromere length = 1 Mb. (B) Centromere length = 2 Mb. (C) Centromere length = 3 Mb.

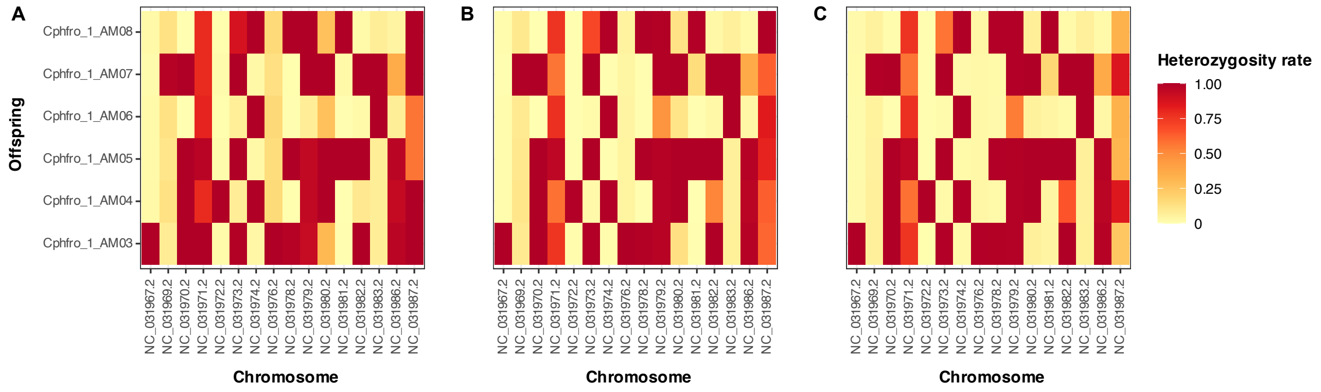
